# Identification of binding interaction between *Curcumin derivative* and *PERK13 Proline rich receptor like protein kinase* protein using *In silico* docking techniques – to help plants tackle salt water

**DOI:** 10.1101/2021.10.15.464501

**Authors:** Prakash Vaithyanathan

## Abstract

**Introduction:** *Arabidopsis thaliana*, mouse ear cress or thale cress are small flowering plants included in the cruciferae family. They comprise of various characteristics such as diploid genetics, small genome size, rapid growth cycle, and relatively low repetitive DNA content, making it a perfect model for plant genome projects.

**Objective:** The aim of this present *Insilico* research study is to carry out molecular drug docking studies between PERK13-Proline rich receptor like protein kinase and De O Acetylated Curcumin Di Galactose or di–galactosylated curcumin, a derivative of curcumin. PERK13 protein is considered to play a significant role in helping plants tackle salinity levels in water.

**Methods:** In this study, protein modelling tools and servers are used to model the 3D structure and the same is validated using Protein structure validation tools. Automated drug docking servers were used to dock the modelled protein with the chemical compound to analyse the electrostatic (H bond) interaction between PERK13 and De O Acetylated Curcumin Di Galactose, a better water-soluble compound than curcumin. The docked structure was visualized using an advanced molecular visualization tool.

**Results and Discussion:** The overall results obtained from this study on De O Acetylated Curcumin Di Galactose and PERK13 protein shows that De O Acetylated Curcumin Di Galactose, directly binds with the active site and other potentially binding regions of PERK13. Hence, it is concluded that De O Acetylated Curcumin Di Galactose could potentially play a vital role in future research related to the problem of helping the plants tackle increased saline levels in water.

## Introduction

*Arabidopsis* model has been adopted world-wide by plant biologists to carry out plant based research^1,2,3,4,5,6,7,8,9^. The root-specific AtPERK13 or PERK13 protein is believed to be playing a significant role to help transport sodium ions from water.^10^

## Methodology

The amino acid sequence of PERK13 (Proline-rich receptor-like protein kinase) protein is shown in Fig. 1a. PERK13 protein was retrieved in FASTA format using UniProt database. The total length of the protein was 710 aa and Mass (Da):75,372 (Fig. 1a). In first step involving protein analysis, we use UniProt Prosite scan database which clearly shows that the amino acids in the range 353 – 619 falls within the functional domains (protein kinases) of PERK13. Protein kinases^11-14^ are enzymes that makeup a large family of proteins that share sequence homology. Their conserved catalytic core is common to serine/threonine as well as to tyrosine protein kinases. There are several conserved regions present in the catalytic domain of protein kinases. Two of these regions were selected to construct signature patterns. The first region, which can be spotted in the N-terminal end of the catalytic domain, is a stretch of residues, which is rich in glycine, and which is present near a lysine residue. It has also been indicated to be concerned with ATP binding (Fig. 1b). The coloured region in Fig. 1b. indicates the various functional domains present in PERK13. In the second step, mobiDB Database (https://mobidb.bio.unipd.it/Q9CAL8) was used for domain analysis. The results obtained from MobiDB Database for PERK13 protein shows that another functional domain of PERK13 lies within 676-700 polar residues (Fig. 1c). The blue coloured region in Fig. 1c represents predominant functional domains of PERK13 (356-626aa). SWISS-MODEL was used to convert the amino acid sequence of PERK13 into 3D structure Fig (1d) and also to elaborately analyse the molecular and structural details of PERK13 for the purpose of docking^15^. The 3D structure of PERK13 viewed in “Solid Ribbon Model” using Discovery Studio Software is shown in Fig. 1d.

**Fig. 1.**
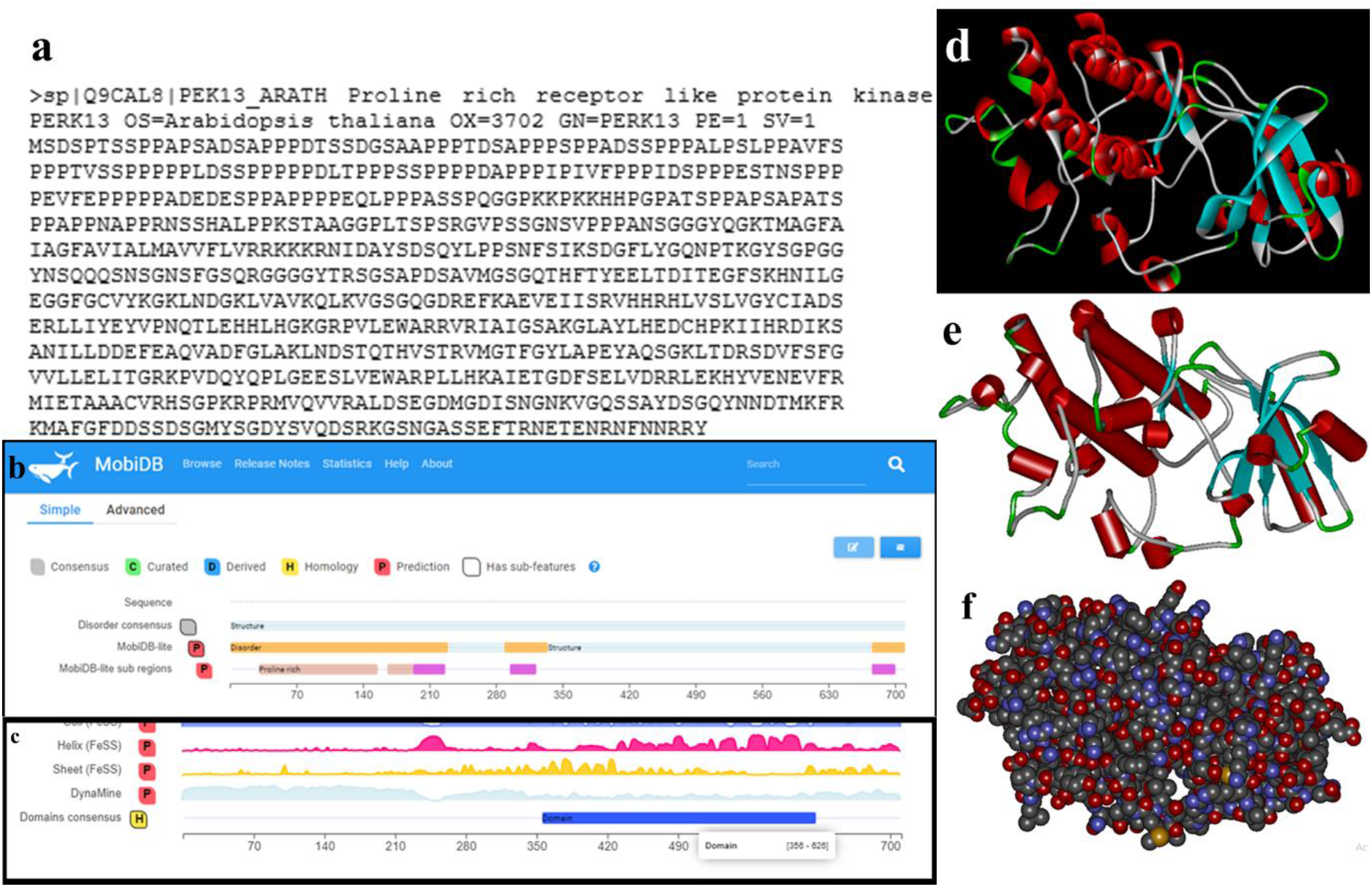
**a**. FASTA sequence of PERK13. b. Sequence analysis of PERK13 using MobiDB Database. c. Sequence analysis of PERK13 using MobiDB Database. d. 3D structure visualization of PERK13 (Solid Ribbon Model). e. Visualization of the 3D structure of PERK13 (Cartoon Model) f. 3D structure visualization of PERK13

Waterhouse et al.^16^ computed models by the SWISS-MODEL server homology modelling pipeline which relies on ProMod3, an in-house comparative modelling engine based on Open Structure.^17^ The 3D structure of PERK13 viewed in “Cartoon Model” using Discovery Studio Software is shown in Fig. 1e and the corresponding 3D structure of PERK13 is shown in Fig. 1f.

The modelled 3D protein was completely evaluated using ProCheck server for assessment of Ramachandran Plot. The results for PERK13 protein obtained from ProCheck server show a Ramachandran plot score of 87.1% which indicates that the modelled structure is acceptable. The assessment of Ramachandran Plot for the modelled PERK13 protein is shown in Fig. 2a and the corresponding statistical analysis is shown in Table 1.

**Fig. 2.**
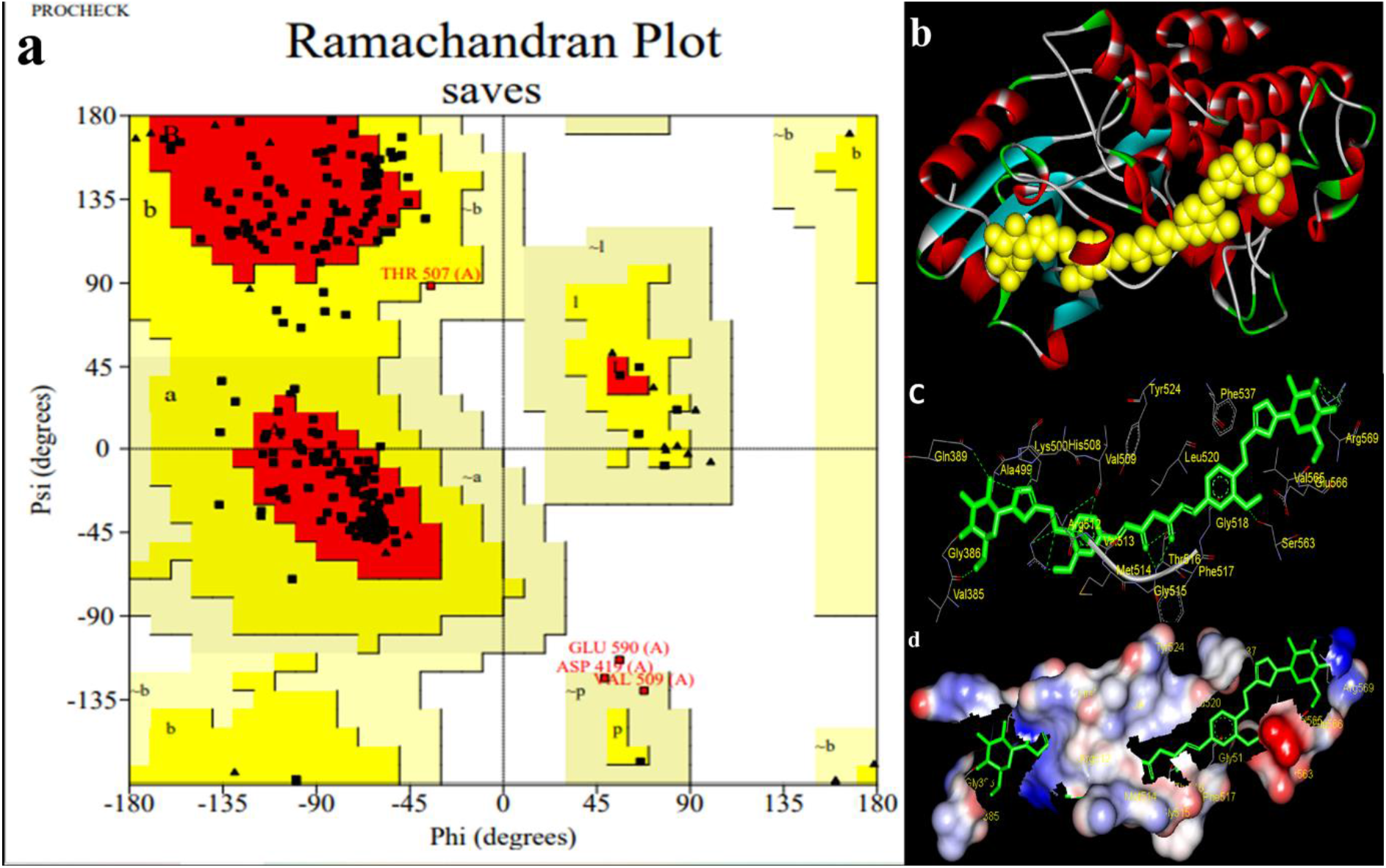
a. 3D structure validation- ProCheck server. b. Molecular drug docking (Digalactosylated curcumin with PERK13). c&d. Prediction of Drug-Protein binding sites in the complex.

**Table 1.** Statistical assessment of Ramachandran Plot for the modelled PERK13 protein using ProCheck server

| Plot statistics |  |  |
| --- | --- | --- |
| Residues in most favoured regions [A,B,L] | 222 | 87.1% |
| Residues in additional allowed regions [a,b,l,p] | 29 | 11.4% |
| Residues in generously allowed regions [~a,~b,~l,~p] | 3 | 1.2% |
| Residues in disallowed regions | 1 | 0.4% |
|  | ---- | ----- |
| Number of non-glycine and non-proline residues | 255 | 100.0% |
| Number of end-residues (excl. Gly and Pro) | 2 |  |
| Number of glycine residues (shown as triangles) | 24 |  |
| Number of proline residues | 9 |  |
|  | ---- |  |
| Total number of residues | 290 |  |
| Based on an analysis of 118 structures of resolution of at least 2.0 Angstroms and R-factor no greater than 20%, a good quality model would be expected to have over 90% in the most favoured regions. |  |  |

The modelled 3D structure of the complex between digalactosylated curcumin and PERK13 gene as viewed using Discovery Studio Software reveals the atomic level intereactions between the water soluble curcumin derivative and the PERK13 gene (Figs 2b, 2c and 2d). The yellow coloured spacefill model represents De-O-Acetylated Curcumin Di-Galactose which binds with PERK13 and is viewed using Discovery Studio Software. 3D structure of the De-O-Acetylated Curcumin Di-Galactose - PERK13 complex is shown in Fig. 2c. Green coloured structure in Stick model represents De-O-Acetylated Curcumin Di-Galactose which interacts with PERK13 at the amino acid binding regions viewed using Discovery Studio Software. The 3D form of the De-O-Acetylated Curcumin Di-Galactose - PERK13 complex is shown in Fig. 2d. Green coloured structure in Stick model represents De-O-Acetylated Curcumin Di-Galactose which interacts with PERK13 (Surface model) at the amino acid binding regions viewed using Discovery Studio Software.

## Results

The selected test compound, *De-O-Acetylated Curcumin Di-Galactose* or digalactosylated curcumin, for short, was docked with PERK13 and the molecular H-bond interaction between them was analysed using Patchdock server.^18, 19^ **Table 2** shows the drug-docking result with Atomic Contact Energy (ACE) value. The PatchDock result scores of PERK13 with *De-O-Acetylated Curcumin Di-Galactose* and the 3D structure of H-bond interactions with the respective drug-binding amino acid pockets (VAL:385, GLY:388, GLY:386, GLN:389, LYS:500, LYS:500, THR:505, ALA:499, HIS:508, ARG:512, VAL:509, ARG:512, ARG:512, ARG:512, ARG:512, GLY:363) were shown in (Tables 2 and 3; Figs 2c and 2d). The best ACE value of De-O-Acetylated Curcumin Di-Galactose with PERK13 protein complex is found to be (−431.21 kcal/mol).

**Table 2.**
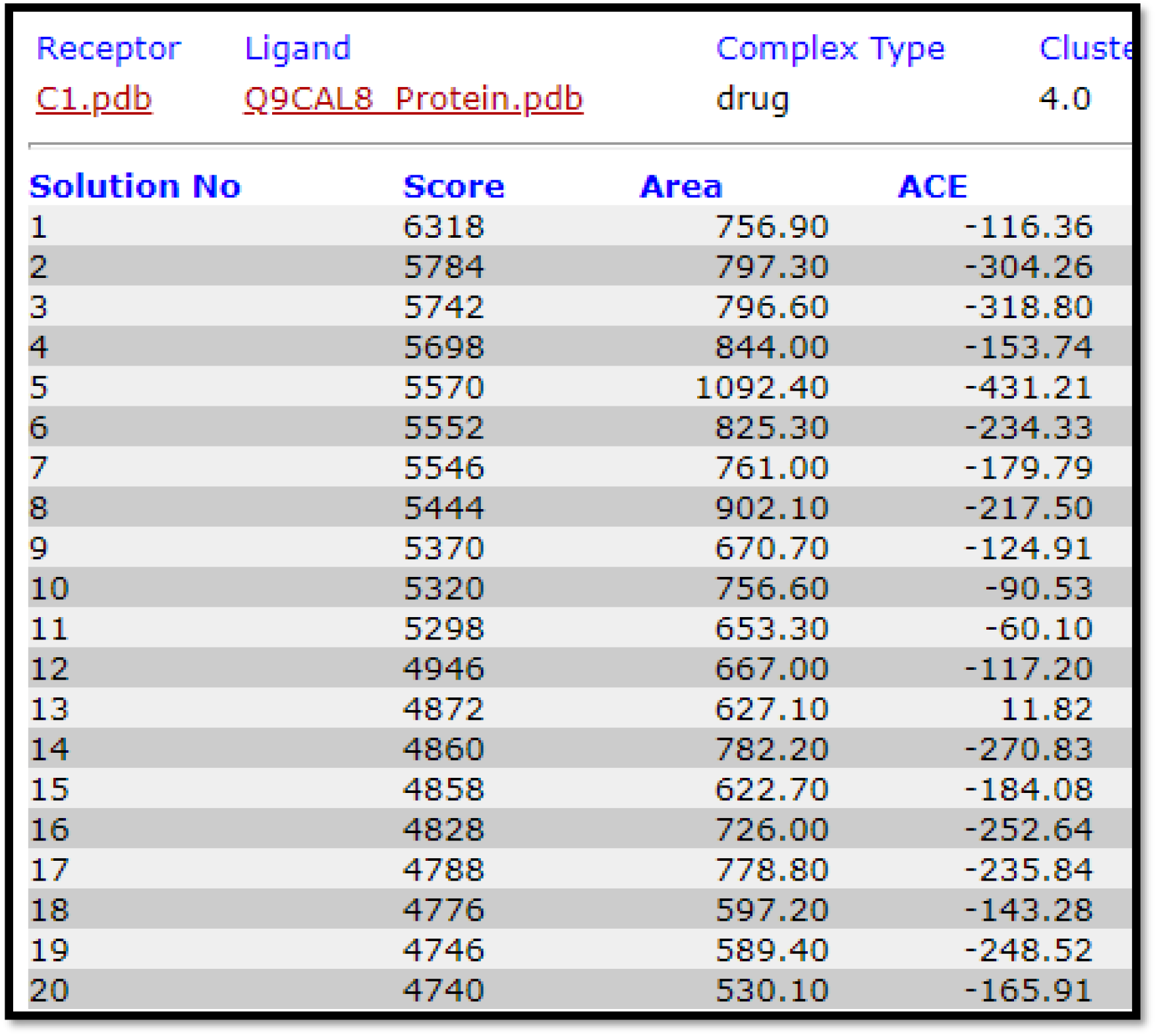
Molecular drug docking – PatchDock results

The Electrostatic and H-bond interaction between De-O-Acetylated Curcumin Di-Galactose and PERK13 are shown in Fig. 3a. The donor with amino acid lables (Fig. 3b) and the acceptor with amino acid labels (Fig. 3c) are also shown. The hydrophilic interactions between the drug and the protein are depicted in Fig. 3d. In Fig. 3a, the left side column indicates the donor amino acids and the right side column indicates the acceptor amino acids The PatchDock results of *De-O-Acetylated Curcumin Di-Galactose* with PERK13 show an ACE (Atomic Contact Energy) value of −431.2 kcal/mol. Higher negative value indicates more binding affinities between the target and the compound. Here, it has been proved that *De-O-Acetylated Curcumin Di-Galactose* is an ideal candidate to interact with PERK13. The ACE (Atomic Contact Energy) value deduced from molecular docking studies is −431.21 kcal/mol.

**Fig. 3.**
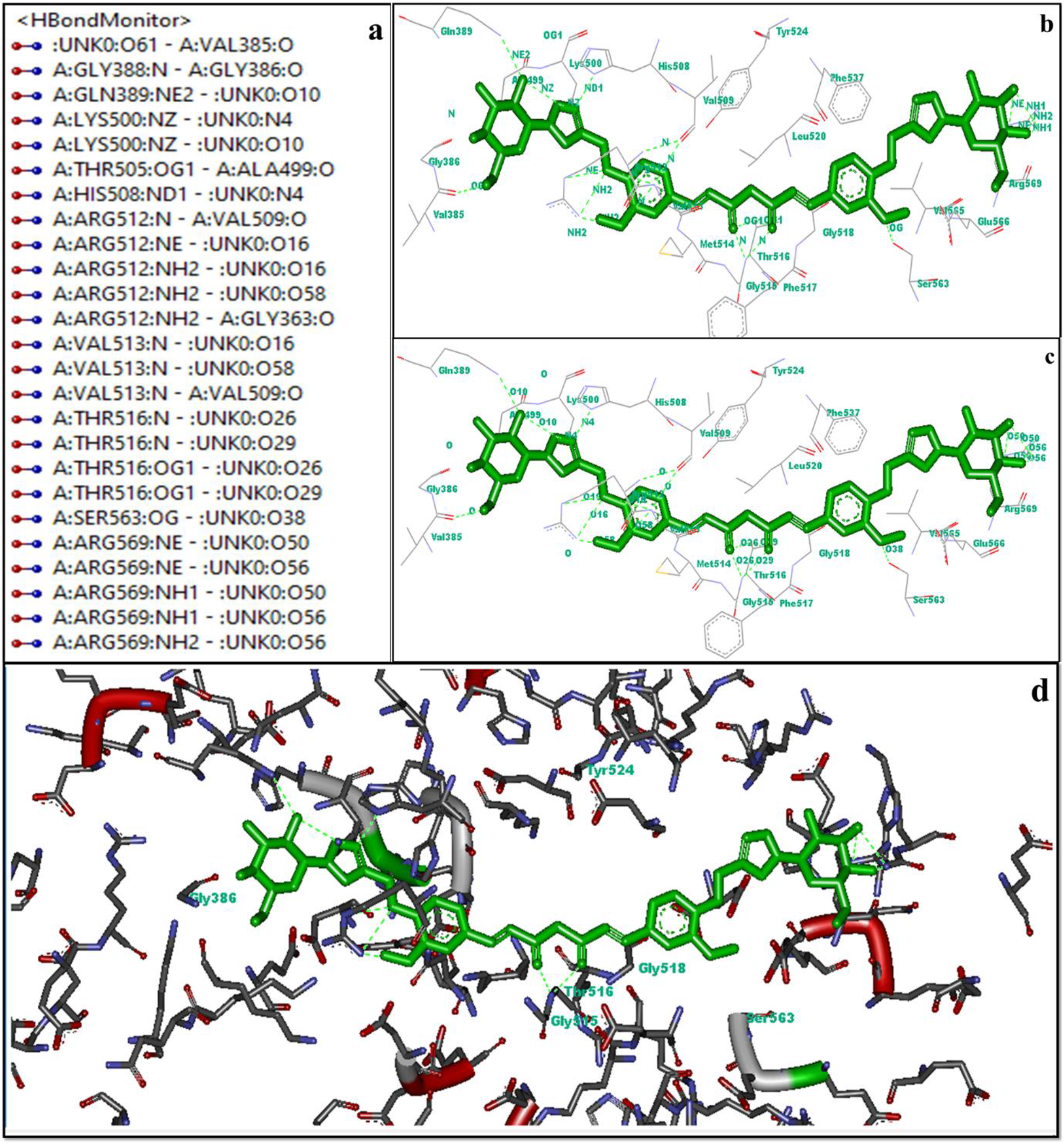
a. H bonding interaction of De-O-Acetylated Curcumin Di Galactose with PERK13. b. Donar with amino acid labels. c. Acceptor with amino acid labels. d. Hydrophilic interaction between the protein and ligand.

When compared with the ACE values of interaction between the PERK13 protein and Curcumin (−221.73 kcal/mol), Neotame (artificial sweetener −275.72 kcal/mol), pyrroloquinoline quinone (−170.33 kcal/mol), the water-soluble test compound used in this study, digalactosylated curcumin^20^ has higher binding affinity (−431.21 kcal/mol).

The chemical structure of the water-soluble curcumin derivate, namely, digalactosylated curcumin, used in this study is shown in Fig. 4a.^20^ Like wise the 3D form of the complex of the drug (water soluble curcumin derivative) and the protein (PERK13) is shown in Fig. 4b with the bond length values depicted in Å.

**Fig. 4.**
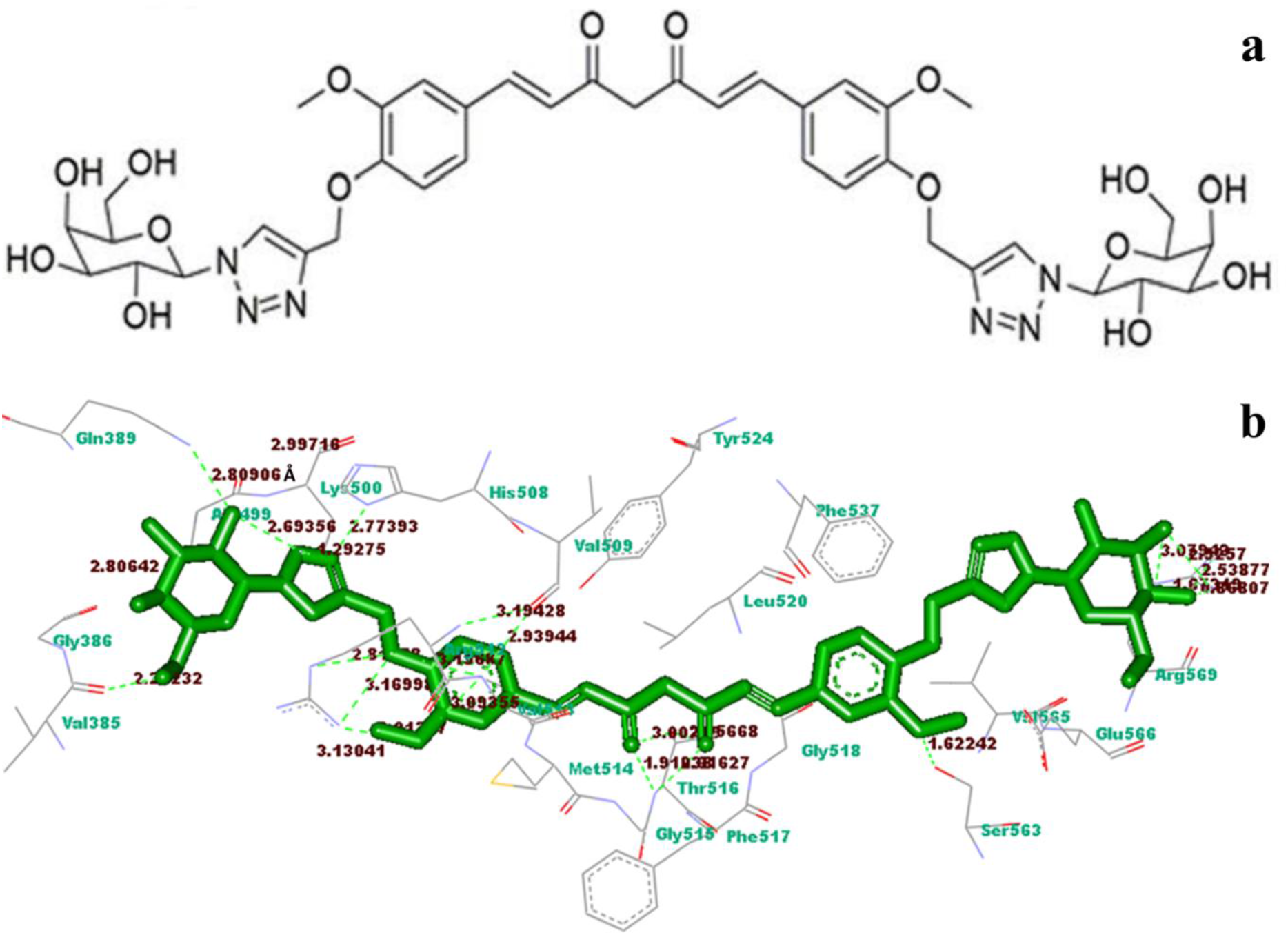
a. Chemical structure of digalactosylated curcumin. b. 3D structure of the complex between digalactosylated curcumin and PERK 13 with specification of the bond length values in Å

Secondly, a modelled structure of the PERK13 protein is retrieved from EBI-Alphafold server and its docking interaction with the ligand, digalactosylated curcumin was studied using patchdock server. The ligand protein interactions using the molecular operating environment tool^22^ demonstrates further the existence of strong pi-pi bond interaction between the ligand and the protein as shown in Fig.5 below.

**Fig. 5.**
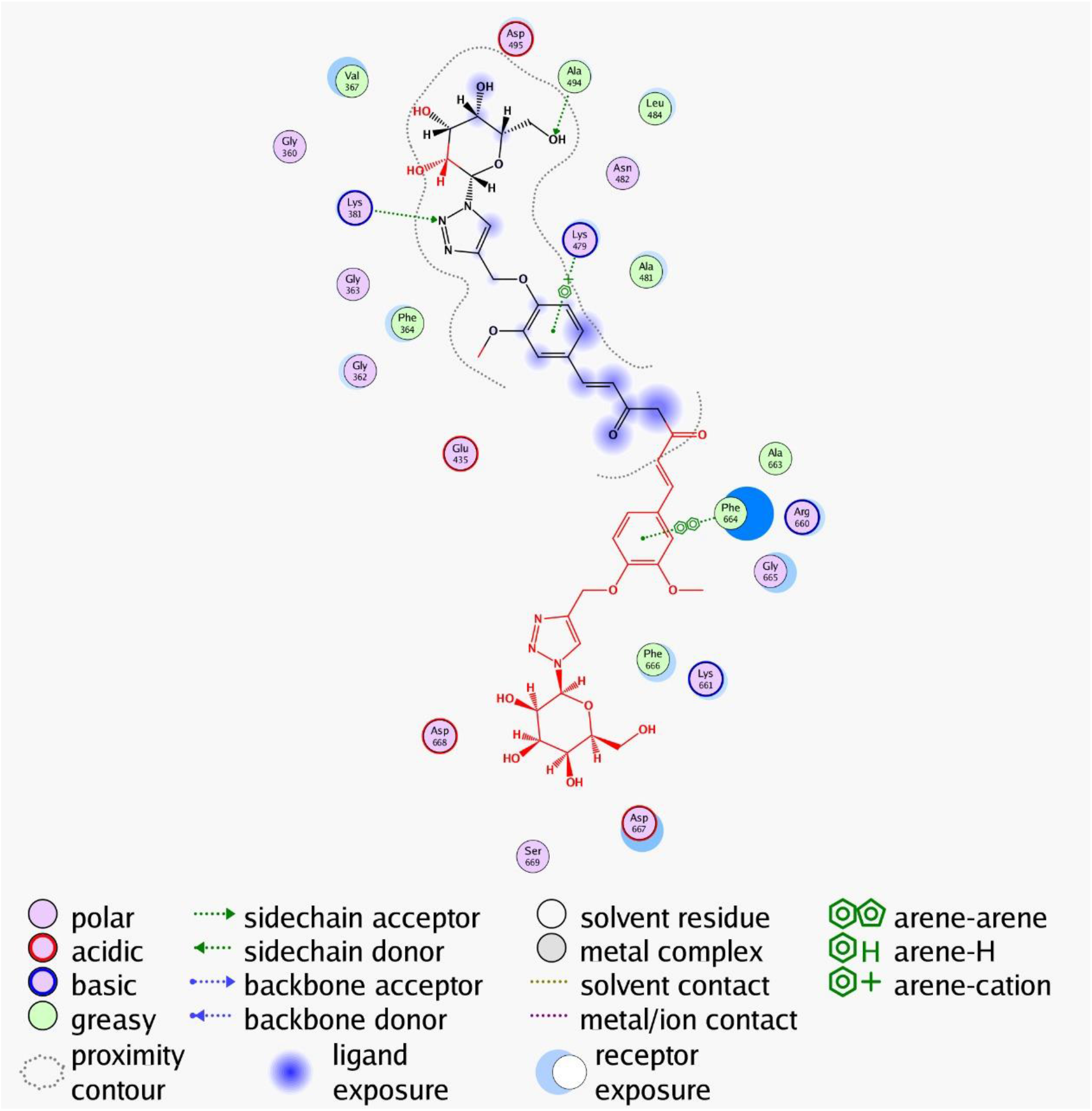
Digalactosylated curcumin and PERK13 interaction

The image demonstrates the role of curcumin and hence the water soluble derivative will be helpful to modulate the functions of PERK13 protein.

## Discussion

The current docking study clearly proves that the Domains and the polar residue amino acid positions (Drug Binding Pockets) of PERK13 exhibit H-bond interaction with De-O-Acetylated Curcumin Di-Galactose and hence shows the potential of the water soluble curcumin derivative^20^ to have the ability to bind strongly to the plant root specific PERK13 protein with application in modulating the salt tolerance of plants. High binding affinity of the test compound can be partially attributed to its hydrophilic wings. This gives a clue to use the hydrophilic signature of the PERK13 protein as a guide to design stronger binding ligands to help plants tackle saline levels in water. It appears by nature, this protein has evolved hydrophilic residues to make it better suitable for its interaction with an aqueous environment.

Hence, the selected compound can be used to check for its antagonistic properties in future studies to help plants tolerate saline levels. Interestingly, PERK13 is also involved in phosphate induced root hair elongation^21^, implying the possibility of using digalactosylated curcumin to study the effects.

## Conclusion

In this current *Insilico* drug docking research study, the selected compound, *De O Acetylated Curcumin Di Galactose* efficiently binds with the target protein, PERK13 (Proline rich receptor like protein kinase). It is believed that the influx of sodium ions through the roots is regulated by PERK13. Hence, the current study will lay the ground for further research to study and solve the problem of salt tolerance of plants. The current docking study clearly proves that the Domains and the polar residue amino acid positions (Drug Binding Pockets) of PERK13 exhibit H bond interaction with *De O Acetylated Curcumin Di Galactose*. Hence, the selected compound can be tried as an efficient agent in future studies to modulate the function of PERK13 gene.

## Acknowledgement

Thanks to the late Sri V Balasubramanian for his help over the years and his son Mr. B V Narayanan (currently residing in U.K.) for funding this project. I would like to extend my thanks to Dr. Balaji Munivelan, Ph.D. for providing paid access to commercial docking software facilities. I thank my student Dr. Harishchander Anandaram, Amrita Vishwa Vidyapeetham for all the basic lessons in bioinformatics.

## Notes

### Summary of Updates

The latest predicted structure of the protein obtained from the AlphaFold protein structure database was used to study the interactions with the ligand and the results are given.

